# Drivers of space use in a large semi-urban feral ungulate under seasonally fluctuating resource conditions

**DOI:** 10.64898/2026.06.03.729827

**Authors:** Debottam Bhattacharjee, Kate J. Flay, Hannah S. Mumby, Jie Zhang, Jin Wu, Alan G. McElligott

## Abstract

Resource scarcity prompts animals to adjust their space use in ways that enhance their survival. In wild herbivores, seasonal habitat shifts are well studied; however, little is known about how large herbivores navigate human-dominated landscapes under fluctuating resource conditions. In Hong Kong, feral water buffalo (*Bubalus bubalis*; henceforth buffalo) experience declines in body condition score during the dry season. While buffalo expand spatial ranges in the dry season, whether this expansion reflects access to improved ecological conditions, and how key anthropogenic factors (like distance to roads and human density) further influence space use, remains understudied. We observed six buffalo herds (*n=92* known individuals) across one wet (July-September 2023) and one dry (January-March 2024) season. We recorded herd locations and extracted remotely sensed habitat and environmental data. We used Normalized Difference Vegetation Index (NDVI) as a proxy of vegetation productivity and included elevation, distance to roads, and human density as additional predictors. We hypothesized that dry-season space use would shift toward higher vegetation productivity, and that anthropogenic factors would have weak influence on space use due to the adaption of buffalo in human-dominated landscapes. We found that buffalo used areas with higher vegetation productivity and higher elevation in the dry season than in the wet. Distance to roads and human density had no detectable effect. Our findings reveal that space use by a large herbivore like buffalo in human-dominated landscapes is strategic and resource driven, and that these seasonal shifts may have important implications for local biodiversity and human–animal interactions.

## Introduction

Understanding how animals adaptively adjust their space use patterns in response to environmental variability is central to behavioral ecology and conservation biology (Aarts et al., 2008; Beever et al., 2017; Kernohan et al., 2001; Owen-Smith, 2008). Selection of habitat and associated movement decisions can directly determine access to forage, water, and resting sites, thus influencing survival (Kernohan et al., 2001; Mueller & Fagan, 2008; Nathan, 2008; Owen-Smith, 2008). Large herbivores are of particular interest because, despite the considerable energetic demands associated with body size, they offset these costs via optimal foraging and energy maximization (Fortin et al., 2024; Owen-Smith et al., 2020). In other words, fluctuations in resource availability may lead to animals using adaptive strategies, such as shifts in home range size, movement patterns, and habitat selection (Abraham et al., 2022). For instance, often in the resource-scarce dry seasons, an expanded space use can buffer individuals against eco-physiological stress (Aikens et al., 2020; Pokharel et al., 2017; Young et al., 2009). Our knowledge of space use dynamics and their drivers in large herbivores predominantly comes from research in the wild. Consequently, little is known about how animals living in complex human-dominated landscapes cope with fluctuating resource conditions and what factors shape their movement and space use decisions.

Human-dominated landscapes are rapidly expanding in the Anthropocene (Lewis & Maslin, 2015), and they present animals with both opportunities and challenges (Linnell et al., 2020; Nyhus, 2016; Pooley et al., 2021; Yirga et al., 2026). While some animals living close to humans can more readily access human-subsidized food resources without the need for costly foraging and may face reduced predation risks, they can also be subject to challenges associated with increased habitat fragmentation pressure and human-animal conflicts (Bhattacharjee & Bhadra, 2021; Gill et al., 2025; Goddard et al., 2010; Oro et al., 2013; Sih, 2013; Wong & Candolin, 2015). Such a complex interplay of anthropogenic factors has implications for the spatial and social ecology of animals (Gaynor et al., 2024). Furthermore, the challenges may intensify in these landscapes if animals, especially large herbivores, face climatic shifts in the availability of natural food resources (Hetem et al., 2014). Therefore, a multitude of factors, from environmental (e.g., seasonal vegetation productivity) to anthropogenic (e.g., human made infrastructure and population density), may drive the behavioral ecology of large herbivores in human-dominated landscapes, and a holistic approach is critical to understanding these mechanisms comprehensively.

In herbivores, vegetation productivity is considered a key determinant of habitat quality and its selection for grazing and browsing (Van Beest et al., 2010). Remote-sensing indices such as the Normalized Difference Vegetation Index (NDVI) are widely used as proxies for primary vegetation productivity or density or greenness, allowing researchers to quantify spatial and temporal variation in plant biomass at ecologically meaningful scales (Borowik et al., 2013). Existing research on wild herbivores have demonstrated that they track spatiotemporal changes in vegetation greenness, often adjusting movements to maximize access to high-NDVI areas (Peters et al., 2017; Seidel & Boyce, 2015). By contrast, access to forage in human-dominated landscapes is often challenging, as these landscapes are characterized by patchy and fragmented habitats, road networks, human settlements, and direct human activities (Goddard et al., 2010; Linnell et al., 2020). Therefore, space use may not only be shaped by natural resource distribution but also by anthropogenic features that may constrain, deter, or, in some cases, attract animals. For instance, roads can act as barriers or increase mortality risks, while human population density may influence disturbance levels, biotic homogenization, supplemental feeding (Bonnot et al., 2013; Eldegard et al., 2012; Mikula et al., 2026; Russo et al., 2026; Yang et al., 2025). Accordingly, assessing how vegetation greenness and features of anthropogenic landscapes jointly influence space use can provide key insights into how large herbivores adapt to human-dominated landscapes.

The semi-urban, feral and free-ranging water buffalo population on Lantau Island, Hong Kong provides a valuable system for investigating the dynamics of space use decisions in large herbivores navigating seasonally variable resources within human-dominated landscapes. The buffalo population inhabits a heterogeneous mosaic of natural vegetation and residential areas in the Southern region of the island (Yang et al., 2025). Previous research has shown that individuals in this population experience declining physiological conditions (*sensu* body condition score) during the resource-scarce dry season in comparison to the resource-rich wet season (Bhattacharjee et al., 2026). Aligning with evidence from other feral buffalo populations that exhibit seasonal shifts in space use in response to changing resource availability (Campbell et al., 2021; Pike et al., 2024, 2026), buffalo on Lantau island also adjust their behavioral strategies during the dry season. Particularly, individuals expand their home ranges beyond the freshwater marshlands and exhibit increased browsing behavior in addition to grazing (Bhattacharjee et al., 2026). However, while such seasonal variation in space use is evident, it does not alone reveal whether movements are strategically directed toward more productive habitats or simply reflect increased searching effort. In other words, a trade-off between energetically costly movement and the potential gains from accessing areas of higher vegetation productivity may shape seasonal space use and movement decisions. Moreover, approximately 10% of Lantau Island’s human population is concentrated in the southern region, which is also characterized by the presence of motorable road networks (Hong Kong Census and Statistics Department, 2024). In recent years, the area has also been under major housing and infrastructural developmental activities, overlapping with the buffalo habitats (WWF-Hong Kong, 2021). Yet, the degree to which these anthropogenic factors influence seasonal space use of buffalo and also potentially their welfare remains unclear.

To investigate the drivers of seasonal space use by feral buffalo inhabiting a human-dominated landscape, we assessed how environmental and anthropogenic factors influence the locations used by buffalo herds across wet and dry seasons on Lantau Island. Given that vegetation productivity strongly influences habitat selection in large herbivores (Aikens et al., 2020; Peters et al., 2017; Van Beest et al., 2010), we first hypothesized that buffalo movements would track spatial variation in forage availability. Specifically, we predicted that during the resource-scarce dry season, when individuals experience declining physiological conditions and expand their home ranges beyond marshlands, buffalo would use areas with vegetation productivity (indexed by NDVI values) comparable to or higher than those used during the wet season. Such a pattern would indicate that seasonal range expansion represents a strategic response aimed at maintaining access to productive forage rather than random increases in searching effort. Second, we examined whether anthropogenic landscape features influence buffalo space use in this human-dominated landscape. As southern Lantau contains scattered settlements and road infrastructure within buffalo habitats, these features may either constrain or facilitate space use behavior. However, given the long-term persistence of buffalo within this landscape and evidence of human tolerance toward buffalo presence (Yang et al., 2025; Zhang, 2025), we predicted that anthropogenic variables, specifically distance to roads and human population density, would have no influence on seasonal space use.

## 2. Methods

### 2.1. Study area and animals

We observed a semi-urban, feral and free-ranging water buffalo population on Lantau Island, Hong Kong Special Administrative Region, China. Buffalo herds on the island are predominantly distributed in its southern region, where they inhabit semi-natural freshwater marshlands interspersed with areas of human settlement (Yang et al., 2025). Observations were conducted on herds present in and around six geographic locations – Luk Tei Tong (Mui Wo) (22°16’06” N 113°59’34” E), Wang Tong (Mui Wo) (22°16’17” N 113°59’46.8” E), Lo Wai Tsuen (Pui O) (22°14’32.7” N 113°58’42.3” E), Lo Uk Tsuen (Pui O) (22°14’29.3” N 113°58’31.1” E), Shap Long Kau Tsuen (22°14’22” N 113°59’27” E), and Shui Hau (22°13’19” N 113°54’48” E) (Yang et al., 2025). As inter-herd interactions were never observed nor reported in this population (Bhattacharjee et al., 2024, 2026), we used the location names to denote the buffalo herds. Across the six locations, the landscape is characterized by a heterogeneous mixture of residential areas and motorable roads, semi-natural freshwater marshlands, slow-flowing streams, backwater rivers, and surrounding grasslands, shrublands, and woodlands that are representative of southern Lantau (**Figure 1**) and broadly of Hong Kong (Dudgeon & Corlett, 2011).

**Figure 1.**
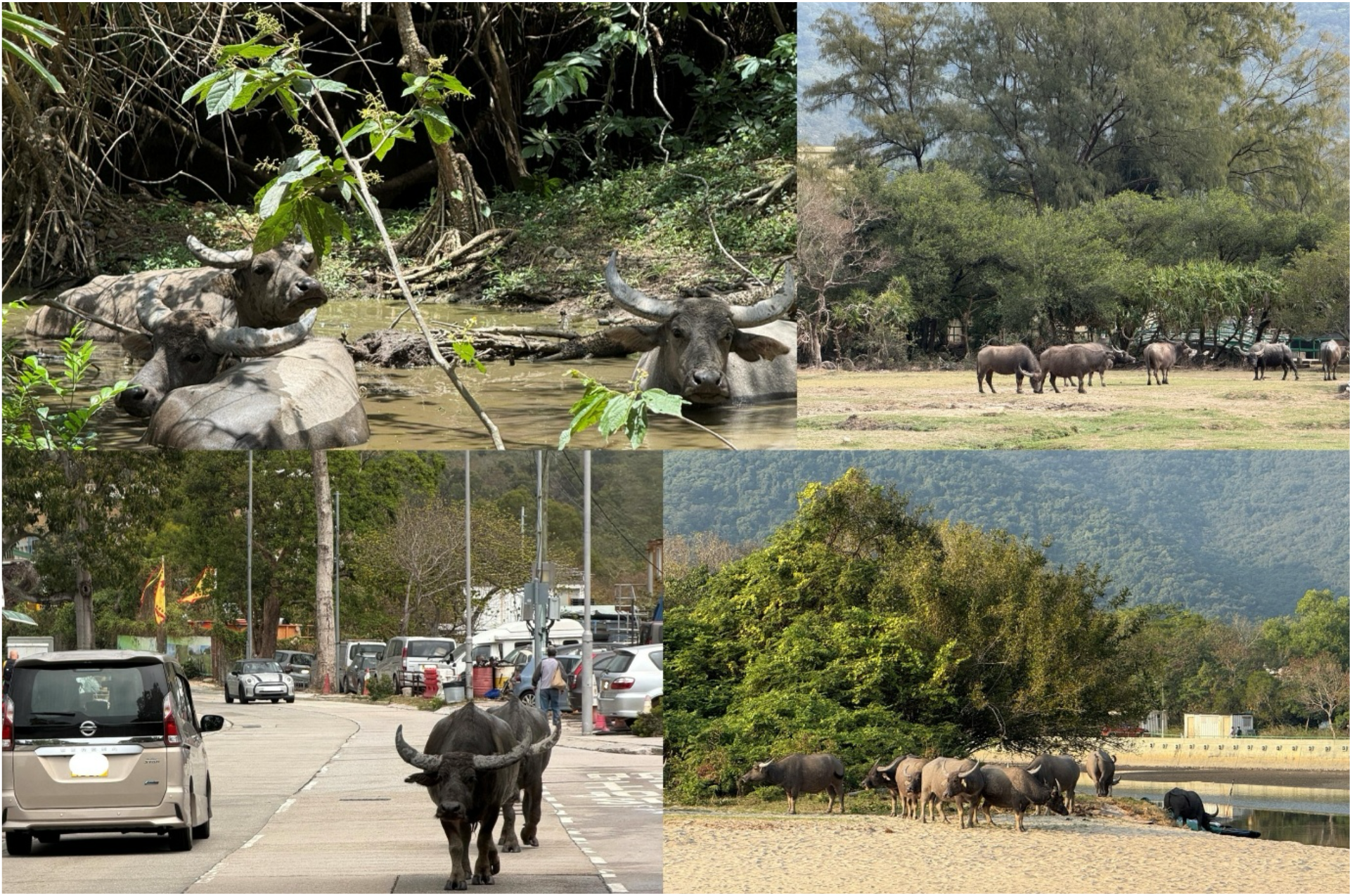
Feral water buffalo in a heterogenous semi-urban habitat on Lantau Island, Hong Kong. Photos showing buffalo using the semi-natural marshlands, grasslands, shrublands and human made infrastructures in Pui O of Lantau Island, Hong Kong. Photo credit: Debottam Bhattacharjee.

We collected data on 92 adult females (age range: 4 – 22 years) belonging to the six herds between July 2023 and March 2024. During the research period, the Lantau Island buffalo population size was 115 (Bhattacharjee et al., 2026, unpubl. Data). Since male buffalo space use, particularly home range size, is known to be restricted by territoriality in this population (Bhattacharjee et al., 2026), we focused only on the females that can move freely across the territories of males (Stammes et al., 2026, unpubl. data). Accordingly, non-herd living solitary males were also not observed. Individual buffalo were identified by their numbered ear tags (administered by the Agriculture, Fisheries and Conservation Department of Hong Kong or AFCD) and our photo catalogue developed based on the morphological features (horn shape and structure, relative horn length, and scar marks) of buffalo. Notably, the Lantau buffalo population occasionally receives supplemental food from local non-government citizen groups, especially during the dry season (Bhattacharjee et al., 2024, 2026; Yang et al., 2025). However, the marked expansion of their home range during the dry season into areas that did not overlap with feeding sites strongly suggests that supplemental feeding had little to no influence on the space use patterns of buffalo (Bhattacharjee et al., 2026).

### 2.2. Ethical note

Non-invasive observations may influence the behavior of animals, thus requiring appropriate ethical approval is necessary (Rowe, 2026). Ethical approval was obtained from the Animal Research Ethics Sub-Committee of City University of Hong Kong (Reference no. AN-STA-00000195). Observations were conducted from a distance of at least 20 m without direct human intervention. We adhered to the ethical guidelines of the ASAB/ABS (ASAB Ethical Committee & ABS Animal Care Committee, 2022). During our non-invasive research, no animals were captured, handled, or manipulated.

### 2.3. Data collection

Data were collected during one wet season (July 2023 – September 2023) and one dry season (January 2024 – March 2024). While both these seasons may span longer than the specified time windows (Bhattacharjee et al., 2026), we avoided collecting data during the transition period, i.e., between October 2023 to December 2023. We took this conservative approach to capture the most climatically distinct periods of the wet and dry seasons, thereby maximizing environmental contrast and its effects. We conducted sampling from 0930 to 1730 hours under daylight conditions. Sampling time was pseudorandomized across locations to ensure representation throughout the daily activity window.

We used a standardized transect sampling method to locate buffalo herds and record their locations by foot. For each location, the transect routes were predetermined (Luk Tei Tong: 3.1 km; Wang Tong: 2.9 km; Lo Wai Tsuen: 2.1 km; Lo Uk Tsuen: 2.3 km; Shap Long Kau Tsuen: 2.5 km; Shui Hau: 3.3 km). Transects followed existing access roads traversing open marshland and grassland habitats where buffalo herds are found with high detectability (cf. Bhattacharjee et al., 2024, 2026). During each transect, the location of a herd was recorded live using a handheld GPS device. Since female buffalo move cohesively as a functional unit, the herd location can provide a biologically meaningful representation of collective space use or movement while avoiding spatial pseudoreplication (cf. (Makarieva et al., 2005)). The herd location was determined where the majority of the individuals (i.e., >50% of respective herds) spatially aggregated. We specifically chose this percentage as it has been shown that group centroid can be estimated reliably with only 50% of group members tagged with GPS devices (He et al., 2023). Nonetheless, on average (± SD), 84.68 ± 11.70% of the herd members were present during data collection. Only one transect per location was conducted on a given day to minimize temporal autocorrelation as buffalo exhibit slow movement and site fidelity (Campbell et al., 2021). We conducted four transects per location per week. From each herd, a total of 45 GPS locations were collected per season (i.e., total 90 GPS locations).

### 2.4. Data preparation and statistical analysis

We performed the statistical analysis using R (version 4.4.1) (R Development Core Team, 2021). For each GPS-recorded herd location, we extracted the environmental and anthropogenic characteristics from publicly available remote sensing and spatial datasets with the help of the following packages: *terra, sf, elevatr*, and *osmdata* (Hijmans, 2020; Hollister, 2025; Maspons et al., 2017; Pebesma, 2018). The variables included vegetation greenness (NDVI derived from Sentinel-2 surface reflectance imagery), elevation (Shuttle Radar Topography Mission or SRTM digital elevation model), distance to roads (OpenStreetMap vector road data), and human population density (WorldPop raster data). Extracting the variables at the herd location coordinates ensured that environmental and anthropogenic conditions corresponded spatially at the time of sampling. All spatial layers were projected to a common coordinate reference system matching the base environmental raster to ensure spatial alignment prior to extraction.

#### 2.4.1. Seasonal NDVI and habitat indices

NDVI was derived from Sentinel-2 Level-2A surface reflectance imagery using Google Earth Engine (GEE). Two seasonal periods were analyzed: wet season (July 2023 to September 2023) and dry season (January 2024 to March 2024). For each period, the *COPERNICUS/S2_SR_HARMONIZED* ImageCollection was filtered by date, spatial extent over Hong Kong, and by scene-level cloud cover (containing <2% cloudy pixels based on metadata). Cloud masking was performed using the Sentinel-2 QA60 band, where bits 10 (cloud) and 11 (cirrus) were required to equal zero. All flagged pixels were removed prior to analysis. Surface reflectance values were then obtained by dividing integer reflectance values by 10,000. NDVI was calculated for each cloud-free image using the standard normalized difference formula:

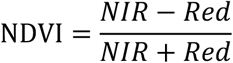

where NIR and Red represent the near-infrared band (Band 8 of Sentinel-2) and red band (Band 4 of Sentinel-2), respectively. The resulting NDVI ImageCollection for each seasonal period was then aggregated into a seasonal mean NDVI composite at 10-m resolution. Finally, the seasonal NDVI composites for both wet and dry seasons were clipped to the study area and exported from GEE in EPSG:4326 for subsequent analysis.

Given the evident spectral mixing between vegetation, exposed substrate, and shallow water in marshland ecosystems as in the case of our study area, we derived additional indices from NDVI to comprehensively represent the habitat structure. Particularly, in marshland-like habitats, NDVI may reflect a composite signal of vegetation density and surface inundation, as water strongly attenuates near-infrared reflectance (Ashok et al., 2021; Ozesmi & Bauer, 2002). As a result, a relatively low NDVI value may correspond to water-flooded marshlands (primarily in wet season where vegetation is submerged under water) instead of a lack of vegetation. Therefore, a spectral wetness proxy was generated by reclassifying NDVI values into ordinal categories representing hydrological and vegetation states. This approach is based on established wetland remote sensing principles, where low or negative NDVI values correspond to open water or saturated substrates due to strong absorption in the near-infrared spectrum, while higher NDVI values represent increasing vegetation productivity or density (Li et al., 2015). The reclassification scheme was based on our empirical NDVI data and defined as: NDVI < 0 (open water / saturated substrate), 0 ≤ NDVI < 0.1 (highly saturated marshland), 0.1 ≤ NDVI < 0.3 (transitional emergent vegetation), and NDVI ≥ 0.3 (vegetated or terrestrial conditions). This ordinal index is hereafter referred to as the NDVI-derived wetness proxy, and is used as a relative measure of habitat inundation gradient rather than a direct hydrological measurement. Furthermore, a vegetation intensity index was derived by truncating NDVI at zero, representing above-ground vegetation greenness while excluding negative values associated with open water or highly reflective surfaces. Finally, a habitat class variable was generated using a modal-rule function within each GPS location, representing dominant habitat state at the spatial scale relevant to buffalo habitat use.

#### 2.4.2. Elevation, distance to roads, and human population density

We obtained topographic elevation data from the SRTM digital elevation model (Jarvis et al., 2004). Elevation tiles covering all herd locations were downloaded at zoom level 12 and converted to a SpatRaster format. We used a bilinear interpolation for the quantification of elevation. Road network data were retrieved from OpenStreetMap (Haklay & Weber, 2008), where we included all road classes (e.g., primary, secondary, and tertiary) to represent human-accessible transport infrastructure. Subsequently, we calculated the Euclidean distances, and for each location, the minimum distance to the nearest road was retained. Human population density was derived from a gridded raster dataset obtained from WorldPop (Stevens et al., 2015). Population density values (persons per grid cell of 0.01 km^2^ from 2025 release) were extracted at each GPS location using raster value extraction.

#### 2.4.3. Statistical models

We fitted linear mixed-effects models (LMM) using the *lme4* and *lmerTest* packages (Bates et al., 2015; Kuznetsova et al., 2017). To quantify seasonal differences in habitat use, we constructed three models reflecting complementary ecological gradients derived from remote sensing. First, the NDVI model, representing overall habitat greenness and integrated vegetation and hydrology signals. Second, a vegetation intensity model that represented positive NDVI values only (NDVI ≥ 0), thus isolating active vegetation structure while reducing potential distortion from open water pixels. Third, a wetness (NDVI-derived wetness proxy) model that represented a reclassified ordinal index of surface inundation and substrate saturation, derived from NDVI thresholds. We fitted three other models where the response variables were elevation, distance to roads, and human population density. All models included season (season: wet / dry) as the fixed effect predictor and herd identity as a random intercept to account for repeated observations within the same social groups. For all full models, we built corresponding null models that lacked the predictor variable, and compared them with the full models using likelihood ratio tests (LRT) from the *lmtest* package (Hothorn et al., 1999). Throughout the statistical analysis, the significance value was set at 0.05.

## 3. Results

### 3.1. NDVI and habitat indices

Seasonal differences in habitat characteristics associated with buffalo herd locations were evident across all indices. The overall NDVI values of locations where herds were observed was higher in the dry season compared to the wet season (LMM: Estimate = 0.117, t-value = 5.99, p < 0.001, **Figure 2a**). A similar pattern was observed for vegetation intensity (i.e., NDVI ≥ 0), which was associated with higher values in the dry than in the wet season (Estimate = 0.113, t-value = 5.89, p < 0.001, **Figure 2b**). By contrast, the NDVI-derived wetness proxy value was significantly lower in the dry season in comparison to the wet season (Estimate = −0.184, t-value = −9.10, p < 0.001, **Figure 2c**). All full models improved significantly from their corresponding null models (NDVI: LRT - χ^2^ = 28.714, p < 0.001; Vegetation intensity - LRT - χ^2^ = 27.628, p < 0.001; Wetness index - LRT - χ^2^ = 71.071, p < 0.001). Together, these complementary results suggest a seasonal shift in habitat use, with buffalo herds using areas characterized by higher vegetation greenness and lower surface wetness during the dry season (**Figure 2d**).

**Figure 2.**
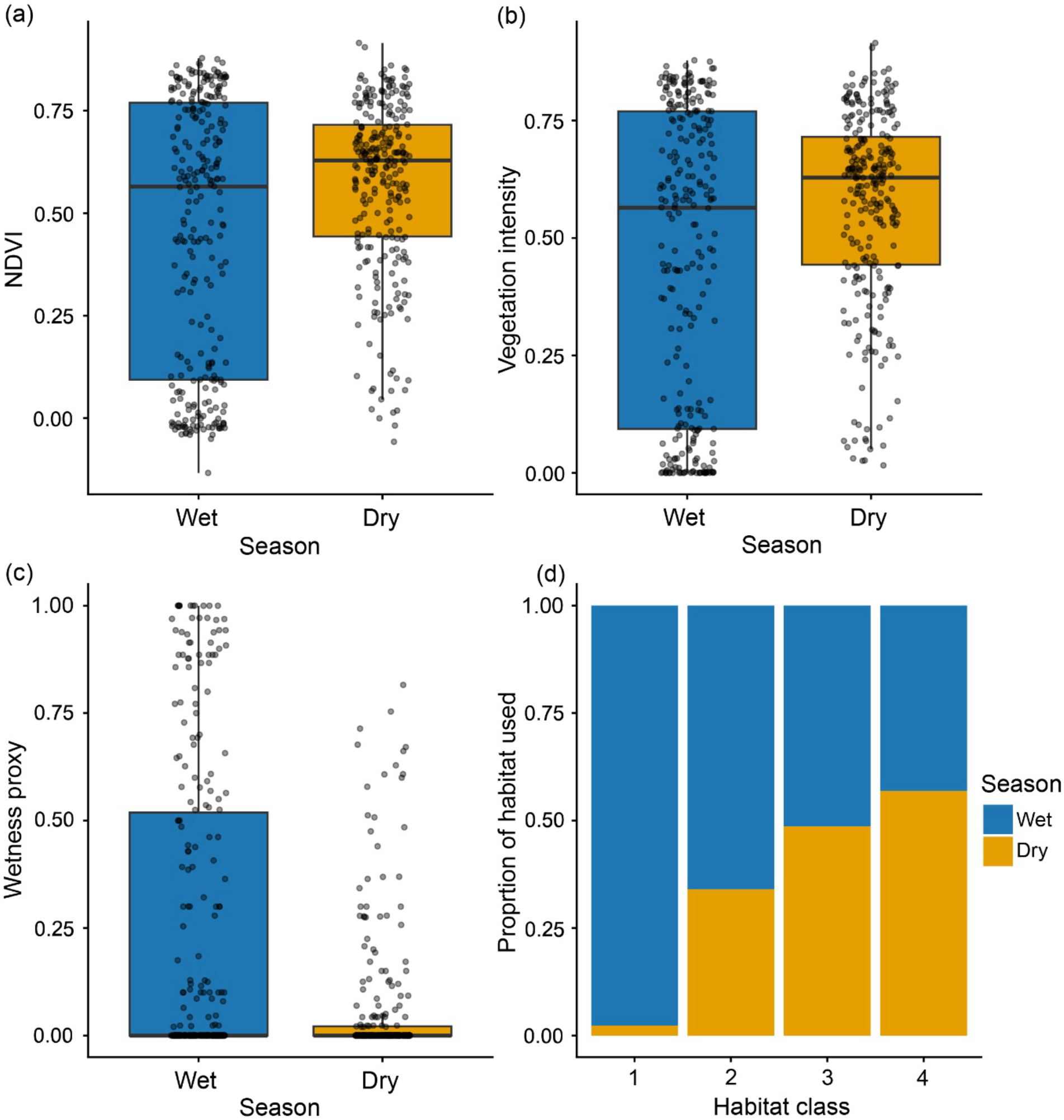
Seasonal variation in habitat characteristics at buffalo herd locations. **(a)** Distribution of NDVI values at buffalo herd locations in wet and dry seasons. **(b)** Vegetation intensity (NDVI ≥ 0) across seasons. **(c)** NDVI-derived wetness proxy values across seasons. **(d)** Proportional distribution of habitat classes derived from NDVI thresholds across seasons. Habitat classes represent a gradient from open water/saturated substrate (Class 1) to dense vegetation (Class 4). The dry season showed a higher proportion of vegetated habitat use by buffalo, whereas the wet season included more water-associated and transitional habitats. In a-c, boxes represent interquartile ranges, and whiskers represent the upper and lower limits of the data. The horizontal bars within the boxes represent median values. Black dots show the distribution of data.

### 3.2. Elevation of used habitat

The mean elevation of areas used by buffalo were 2.37 ± 3.81 m for wet and 9.13 ± 21.65 m for the dry season. Thus, approximately 7 m higher on average areas were used in the dry compared to the wet season. Season had a strong effect on elevation of used areas after accounting for herd identities (LMM: Estimate: 6.76, t-value = 5.19, p < 0.001, **Figure 3**). The full model differed significantly from the null model (LRT: χ^2^ = 28.687, p < 0.001).

**Figure 3.**
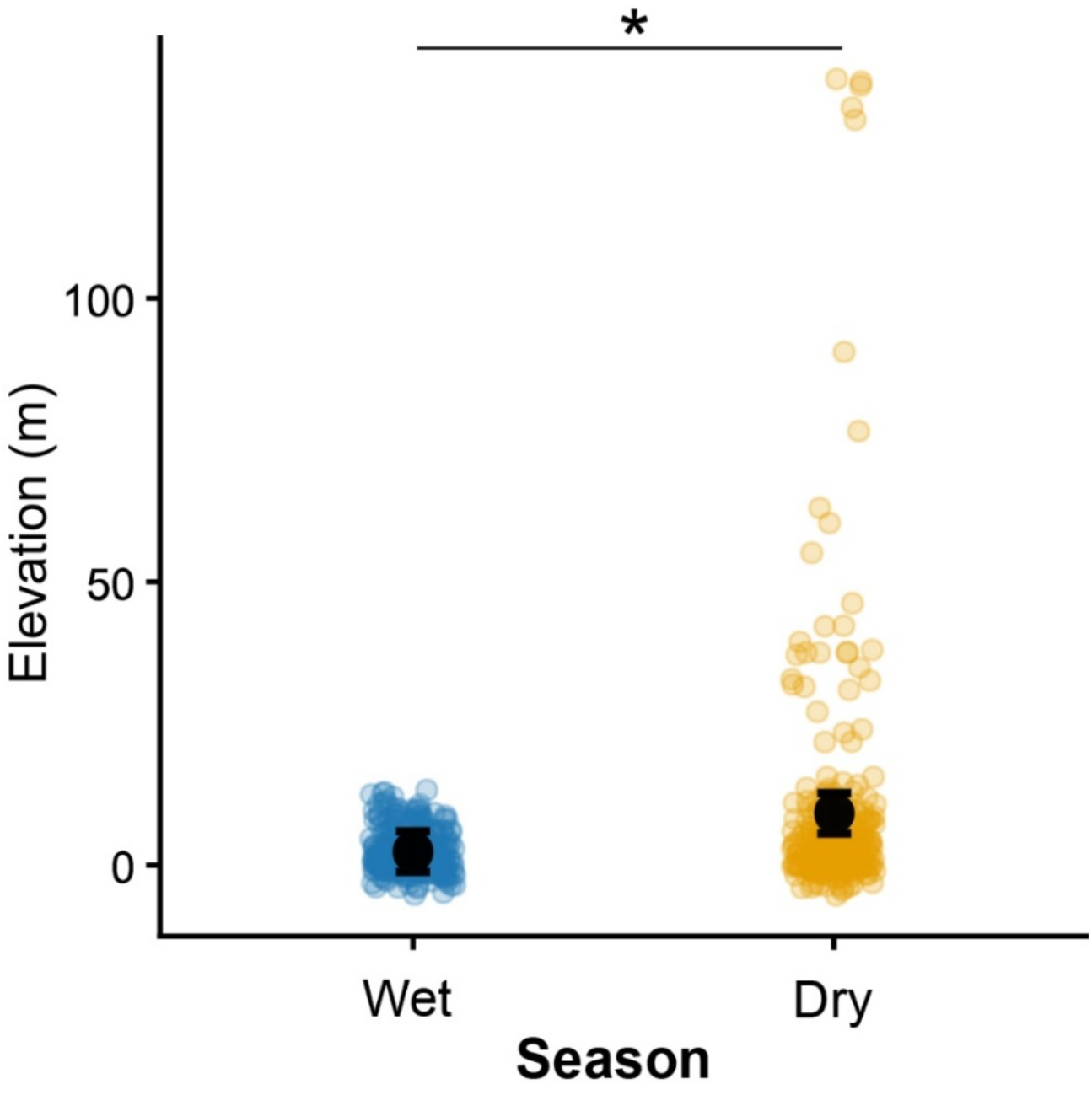
Seasonal pattern in elevation of areas used by buffalo. Solid black points represent model-predicted marginal means from linear mixed-effects models, with 95% confidence intervals (95% CI) shown as black error bars. Faded points show the observed data. Significant effect is highlighted by asterisk.

### 3.3. Distance to roads and human population density

The mean distance to roads were 101.98 ± 55.22 m for the wet and 113.93 ± 123.64 m for the dry season. Seasonal space use patterns were not associated with distance to roads (LMM: Estimate: 11.952, t-value = 1.50, p = 0.13, **Figure 4a**). Human population density of the areas used by buffalo were 2.53 ± 1.84 per 0.01 km^2^ for the wet and 2.49 ± 1.95 per 0.01 km^2^ for the dry season. Similar to distance to roads, we did not find its association with seasonal space use (LMM: Estimate: -0.06, t-value = -0.65, p = 0.51, **Figure 4b**).

**Figure 4.**
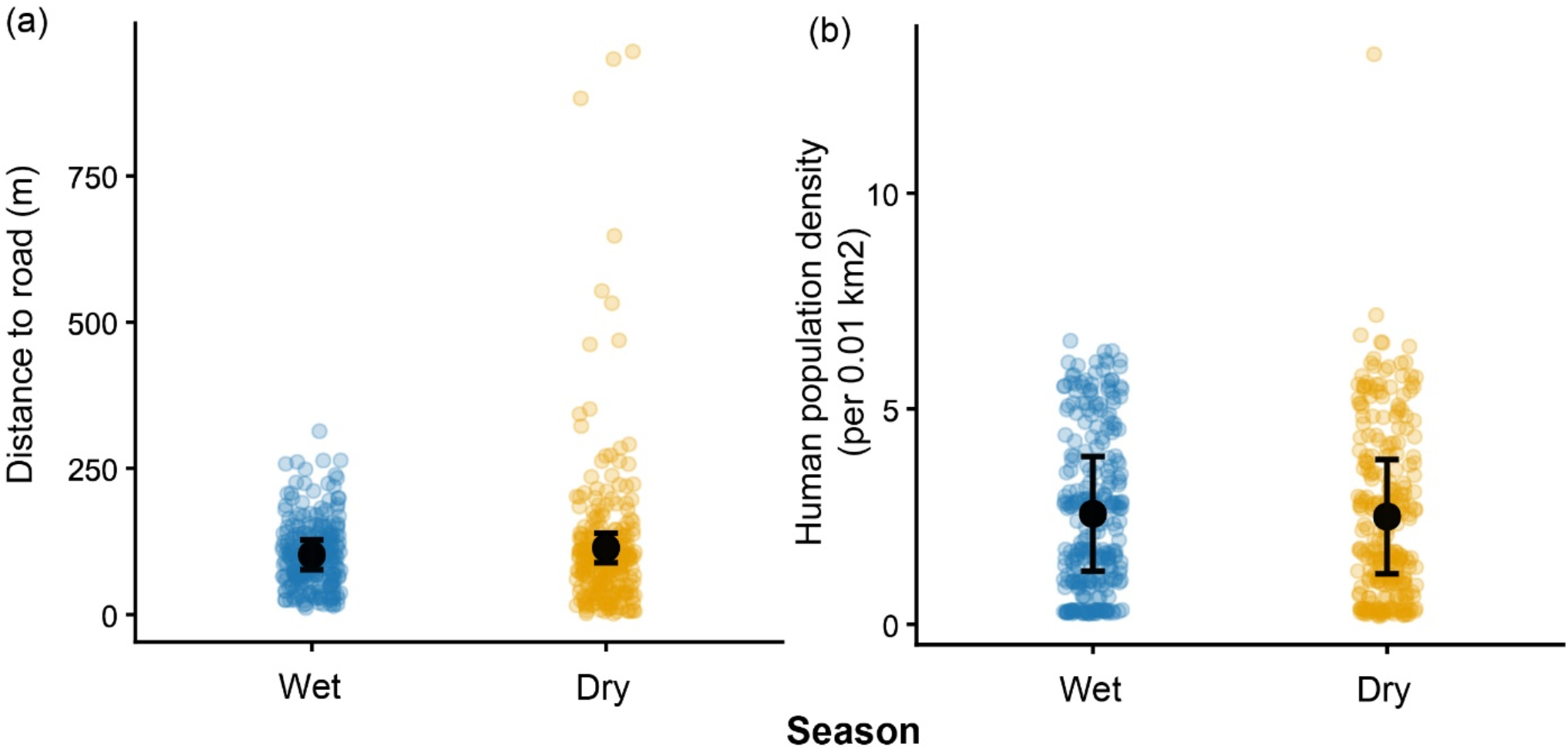
Seasonal patterns in distance to roads and population density of areas used by buffalo. (a) Distance to roads in meters, (b) Human population density per 0.01 km^2^. In both a and b, solid black points represent model-predicted marginal means from linear mixed-effects models, with 95% confidence intervals (95% CI) shown as black error bars. Faded points show the observed data.

### 3.4. Inter-herd differences

Formal statistical testing of herd-level differences was not feasible due to the presence of only six herds. Therefore, herd identity was included as a random effect in all models to account for non-independence of observations within herds and not as random slope. Nonetheless, examination of predicted values (**Figure S1**) revealed inter-herd variations. For instance, predicted overall NDVI values in the dry season ranged from 0.50 to 0.62 across herds, elevation ranged from 5 to 14 m, distance to roads from 90 to 120 m, and population density from 1.5 to 3.2 persons per 0.01 km^2^ (**Figure S1**). These patterns indicate that, in addition to seasonal effects, spatial heterogeneity among herds likely contributes considerable variation in space use.

## Discussion

Adjustment in space use pattern is a central component of large herbivore behavioral ecology that allows individuals to balance energetic demands with spatial and temporal variation in resource availability (Aikens et al., 2020; Fortin et al., 2024; Owen-Smith et al., 2020). In rapidly growing human-dominated landscapes, however, movement decisions may also be shaped by anthropogenic features such as infrastructure, habitat fragmentation, and human disturbance (Gaynor et al., 2024; Linnell et al., 2020). Understanding the influence of environmental resource gradients and anthropogenic landscape elements on animal space use is therefore critical to predicting how large herbivores persist in modified ecosystems. While previous research has shown that feral and free-ranging buffalo expand their home ranges during the resource scarce dry season (Bhattacharjee et al., 2026), the mechanisms underlying such decisions remain poorly understood. We examined how seasonal variation in vegetation productivity and anthropogenic habitat characteristics influence space use by a feral buffalo population inhabiting a semi-urban landscape. We found that at the population level, buffalo visited areas with higher overall NDVI during the dry season than the wet and more frequently used higher elevations. This pattern was consistent also after controlling for the wetness of marshlands associated with NDVI calculations. As per our predictions, proximity to roads and human population density were not associated with space use by buffalo. Beyond advancing our knowledge of large herbivore behavioral adaptation particularly in Asia, our research may have broader implications for local biodiversity, nature of human– feral herbivore interactions in (semi-)urban ecosystems, and for animal welfare (Lundgren et al., 2024; Sih, 2013; Trepel et al., 2026; Yang et al., 2025).

Seasonal space use by buffalo was affected by environmental resource distribution rather than by random exploratory movements across the landscape. During the dry season, buffalo herds visited areas with higher overall NDVI compared to the wet season. The pattern was confirmed after accounting for the wetness of marshland habitats from the NDVI-derived wetness proxy and vegetation indices. This finding supports our prediction that dry season range expansion represents a strategic and adaptive behavioral response aimed at maintaining access to productive forage despite declining physiological condition (Bhattacharjee et al., 2026). For large herbivores, movement decisions are often shaped by the energetic trade-off between costs of movement and foraging benefits (Fortin et al., 2024). Our results suggest that buffalo may resolve this trade-off by directing movements from marshlands toward relatively productive habitats. Similar resource driven patterns have been observed in other feral buffalo populations as well as in other wild ungulates (Calosi et al., 2025; Campbell et al., 2021; Peters et al., 2017; Pike et al., 2024; Seidel & Boyce, 2015), where vegetation productivity (often *sensu* NDVI) is tightly linked to body condition and fitness outcomes, reinforcing the mechanistic importance of vegetation dynamics in shaping space use decisions.

The more frequent use of higher elevation areas in the dry season in comparison to wet further supports the interpretation that buffalo movements are linked to resource dynamics. During the dry season, buffalo used areas approximately 7-m higher in elevation than during the wet season. The use of higher elevation habitats has also been reported in wild ungulates, e.g., in mountain goats, in response to climatic changes (Michaud et al., 2024). Although the absolute elevation differences were not striking given the hilly landscape of the Southern Lantau region, such topographic changes may correspond to variation in vegetation structure and moisture availability. In tropical areas, altitude affects vegetation structure in dry seasons (Gallardo-Cruz et al., 2009), and thus areas with higher elevation may maintain vegetation productivity for longer periods during dry conditions, potentially providing more reliable forage resources. Notably, buffalo generally prefer low-lying coastal marshlands and are naturally associated with habitats close to sea level (Bhattacharjee et al., 2024, 2026; Yang et al., 2025). Therefore, seasonal use of higher elevation areas may not necessarily reflect an intrinsic preference for higher elevations, but instead a temporary displacement from low-lying marshland habitats, potentially driven by seasonal reductions in forage availability. Although not experimentally tested, using higher elevation areas temporarily in the dry season than during the wet season may indicate that buffalo use a “green-wave surfing” rule (*sensu* green wave hypothesis) (Middleton et al., 2018; Ortega et al., 2025).

From a welfare perspective, the seasonal shift in habitat use, including the increased use of higher elevation areas during the dry season, may benefit buffalo by allowing access to more reliable forage resources during a period generally associated with food scarcity for feral bovids in Hong Kong (Bhattacharjee et al., 2026; Perroux et al., 2026). Therefore, dry-season shifts toward greener and higher habitats may not only represent optimal foraging strategies but also behavioral mechanisms supporting resilience. In the context of Lantau Island, the use of elevated areas may also reflect a shift toward more structurally diverse habitats where browsing opportunities increase, i.e., from low-lying food-scarce marshlands to higher elevation shrubs, plants, and grass patches. This finding also aligned with previous findings that buffalo exhibit greater browsing behavior during the dry season (Bhattacharjee et al., 2026). However, we examined space use across one wet and one dry season, which allowed us to capture seasonal contrasts but may not fully represent interannual variability in vegetation dynamics or climatic conditions. Long-term monitoring would therefore help determine whether the patterns observed in our research remain consistent, further providing useful information about welfare of this neglected species (Bukhari et al., 2024). Nonetheless, our results indicate that seasonal space use adjustments likely involve multiple interacting ecological factors, including vegetation productivity and local habitat heterogeneity.

In contrast to environmental drivers, the two anthropogenic landscape features (i.e., distance to roads and human population density) were not associated with buffalo space use. These findings potentially suggest a degree of tolerance and/or behavioral adaptation to human presence in this buffalo population. Animals inhabiting long-established human-dominated landscapes may exhibit reduced avoidance responses if human activity is predictable or relatively non-threatening (Gaynor et al., 2024). Buffalo were introduced to Hong Kong as draught animals and ever since the industrialization during early 1970s, they have been left abandoned, eventually establishing feral populations around human settlements (Cockrill, 1976; Yang et al., 2025; Zhang, 2025). An additional non-mutually exclusive explanation would be that tolerance toward anthropogenic features may be mediated by social information use. In other words, if experienced individuals regularly use areas near roads without exhibiting fear-related behaviours, younger or less experienced herd members may learn that these features do not reliably predict risk, leading to socially transmitted tolerance across generations (Hahn et al., 2026). From a comparative perspective, this pattern contrasts with systems where population density or human disturbance interacts synergistically with environmental stressors to shape condition and behaviour. The absence of such effects in our study may indicate that buffalo are below critical density thresholds or that access to productive habitats buffers potential anthropogenic constraints. Occasional supplemental feeding from local people can also be interpreted as reason for such tolerance, further reducing the perceived risks associated with humans (Bhattacharjee et al., 2024, 2026). Nevertheless, to a small extent, negative human-buffalo interactions have also been reported on Lantau Island (Loand & Westbrook, 2021; Yang et al., 2025). Overall, the anthropogenic features may represent neutral elements of the landscape rather than strong drivers of buffalo space use and habitat shift patterns. Although we considered key anthropogenic variables such as road proximity and human population density, other forms of human activity, such as tourism intensity and direct human-buffalo interactions, may also influence space use and hence should be investigated.

Beyond their own survival strategies, the spatial behavior of buffalo may also have broader ecological and biodiversity consequences. As generalist bulk-feeders, buffalo are physically constrained in their ability to selectively consume specific plant species and therefore tend to graze across a wide range of vegetation types (Lundgren et al., 2024). Such feeding behavior can suppress competitively dominant plant species and create opportunities for less competitive species to persist, thereby increasing overall plant diversity (Lundgren et al., 2024; Trepel et al., 2026). Consequently, seasonal space use by buffalo in productive habitats may shape vegetation structure and plant community composition. In addition to their grazing effects, buffalo are recognized as marshland ecosystem engineers that influence multiple ecological processes. For instance, through activities such as trampling, wallowing, and movement, buffalo can enhance soil nutrient cycling, disperse seeds via dung and fur, facilitate juvenile tree growth, and modify microhabitats that support diverse (semi-)aquatic and terrestrial organisms (Mihailou & Massaro, 2021; So & Dudgeon, 2020; Werner et al., 2006). In this context, the observed seasonal shifts in buffalo space use may contribute to spatial heterogeneity in plant biomass, soil processes, and potentially carbon dynamics within marshland ecosystems. Collectively, these processes suggest that large herbivore like buffalo, even in feral or non-native contexts, can function as important ecosystem engineers with the capacity to influence biodiversity and ecosystem functioning (e.g., rewilding processes) (Kristensen et al., 2026; Lundgren et al., 2024). Importantly, at the same time, the observed inter-herd variation in habitat use suggests that ecological impacts may not be uniform across the population. Over longer timescales, consistent differences in habitat use among herds could contribute to fine-scale population structuring, where habitat shift shapes ecological and evolutionary dynamics (Owen-Smith, 2008; Owen-Smith et al., 2010). Future research should therefore examine whether individual- and group-level factors, such as age, group size and composition, and social associations, predict variation in space use, and an integrative approach of behavior, ecology, and welfare would provide a comprehensive understanding of how animals navigate and thrive human-modified landscapes.

## Data availability

All data generated from this research and necessary codes for analyses will be made available during peer review (for reviewers) and upon publication (for public).

## Author contributions

Conceptualization: D.B., K.J.F., H.M., A.G.M.; Methodology: D.B., J.Z.; Investigation: D.B.; Formal analysis: D.B., J.Z.; Data curation: D.B.; Writing – original draft: D.B.; Writing – review & editing: D.B., K.J.F., H.M., J.Z., J.W., A.G.M; Funding acquisition: K.J.F., H.M., A.G.M.; Resources: J.W., A.G.M.; Supervision: A.G.M.

## Declaration of Interest

The authors have no competing interests.

## Acknowledgments

We thank Jean Leung of the South Lantau Buffalo Society for providing helpful insights into the Lantau buffalo population. This research was funded by the Lantau Conservation Fund (Hong Kong SAR Government-funded programme; Grant ref. RE-2021-01). J.W. was supported by the Innovation and Technology Fund (funding support to State Key Laboratory of Agrobiotechnology). We thank Steeve Côté, Jonathan Farr, George Hodgson, Benjamin Laurie, and Tania Perroux for their helpful suggestions to our research.

## Supporting Information

**Figure S1.**
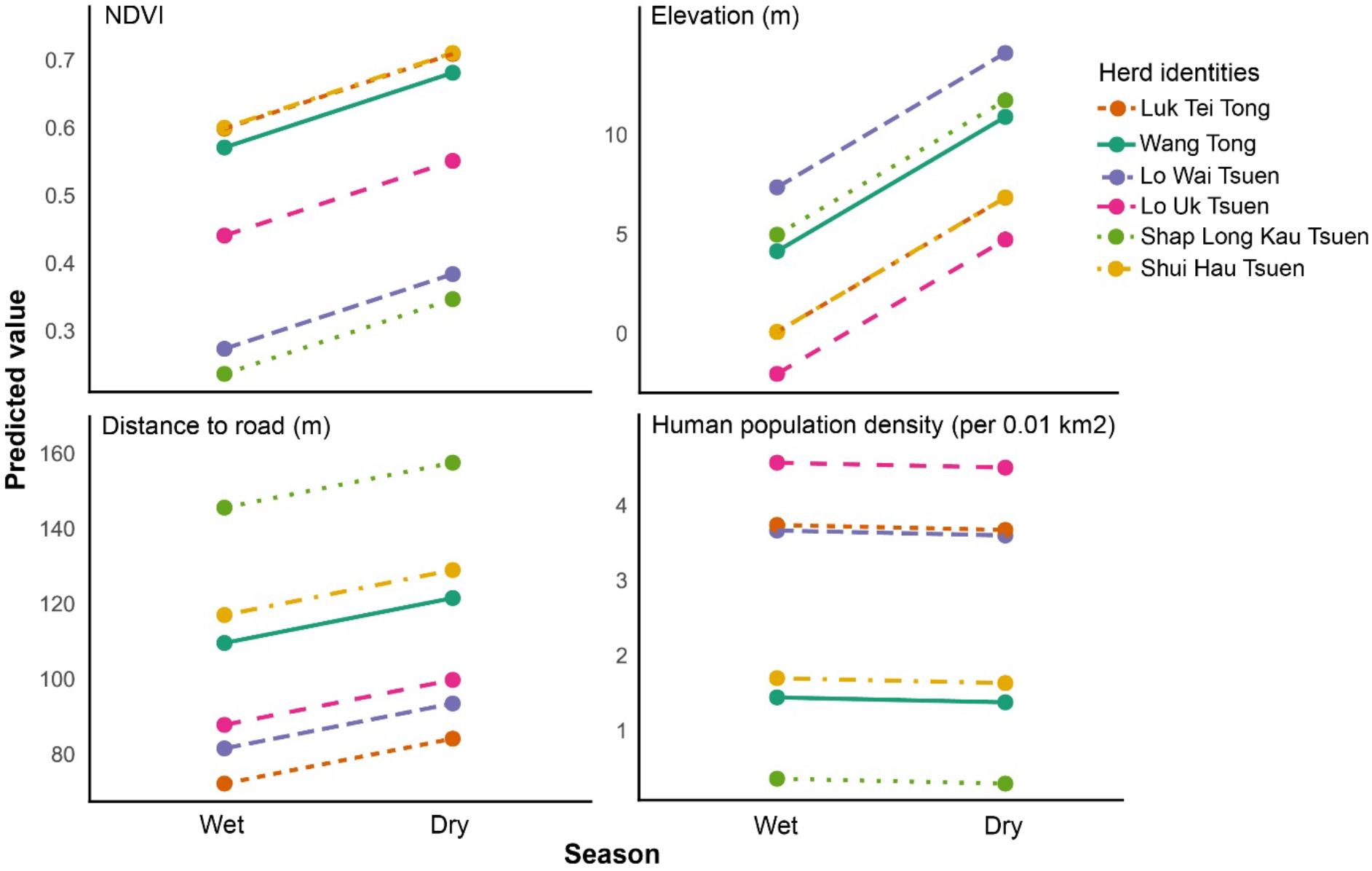
Predicted values of overall NDVI, elevation, distance to road, and human population density across wet and dry seasons for areas used by each buffalo herd.

## Notes

### Competing Interest Statement

The authors have declared no competing interest.

